# 6-Methylnicotine is a strong agonist at α4β2 and α3β4 nicotinic cholinergic receptors

**DOI:** 10.64898/2026.10.09.757661

**Authors:** Thao T. Olson, Reniel Suarez Gonzalez, Yousef Aljohani, Ghazaul Dezfuli, Andrew C. Harris, Mark G. LeSage, Gerard P. Ahern, Kenneth J. Kellar

## Abstract

(-)6-Methylnicotine (6-MN), is a structural analog of (-)nicotine that has been added to some e-cigarette/vape solutions and to nicotine oral pouches. 6-MN binds with high affinity to nicotinic cholinergic receptors (nAChRs) in brain, but its receptor subtype selectivity and its potency to activate and desensitize these receptors is unknown. Also unknown is whether chronic administration of 6-MN increases (upregulates) nAChRs in brain as does nicotine. Currently, 6-MN is unregulated in the U.S. by the FDA. Here we compared 6-MN and nicotine at α4β2 and α3β4 nAChRs, which are two major nAChR subtypes that mediate important functions in the brain and autonomic nervous system. Specifically, we quantitatively compared (-)6-MN and (-)nicotine with respect to receptor binding affinity and functional potency to activate α4β2 and α3β4 nAChRs, as well as its potency to desensitize these receptors, which is one of the key effects of nicotine. In addition, because chronic administration of nicotine upregulates the number of α4β2 nAChRs in rat, mouse and human brain, we measured whether 6-MN also increases these receptors. Our results indicate that 6-MN is a more potent agonist than nicotine at both the α4β2 and α3β4 nAChRs, and it also upregulates nAChRs in brain.

## Introduction

Nicotinic acetylcholine receptors (nAChRs) are ligand-gated cation channels that play essential roles in synaptic signaling critically involved in many functions in the CNS and peripheral nervous systems, including reinforcement and attention processes in the CNS and ganglionic neurotransmission in the autonomic and sensory nervous systems. There are multiple subtypes of these receptors, defined by their subunit composition, which display different biophysical and pharmacological properties, but among these diverse subtypes, the α4β2 and α3β4 receptors are of special interest due to their anatomical distributions and physiological roles. For example, the α4β2 subtype is one of the predominant nAChRs in the brain and is upregulated in rat and mouse brain by chronic administration of nicotine (1, 2) and increased in human brain from people who smoked cigarettes (3, 4). This nAChR is believed to be significantly involved in nicotine addiction and dependence, as well as in reward, cognition, and affect. (5-9). The α3β4 receptors are highly expressed in autonomic ganglia (10) where they mediate the signals for critical autonomic nervous system functions; and they are also highly expressed in the brain’s medial habenula (MHb) and interpeduncular nucleus (IPN), where they are thought to mediate certain responses to nicotine and particularly nicotine withdrawal (11, 12) reviewed in (9). Modifications of nicotine’s structure provide an important approach to better understand the molecular parameters of structure-activity relationships for nAChR pharmacology, including activation, competitive and non-competitive antagonism, and regulation, as well as analysis of nAChR subtype selectivity. (-)6-Methylnicotine (6-MN), a structural analog of (-)nicotine, contains a methyl group at the 6-position of the pyridine ring, a modification that may alter ligand-receptor interactions through steric and/or electronic effects. Such changes have the potential to influence both binding affinity and functional potency at specific nAChR subtypes. Even subtle alterations in the structure of a small molecule such as nicotine could shift its receptor subtype selectivity and efficacy at nAChRs, as well as its subsequent cellular effects, highlighting the sensitivity of nAChRs to chemical modification (13-15). 6-MN is a good example of this. It competes with [^3^H]-nicotine for orthosteric binding sites in rat brain and elicits behavioral responses (hypolocomotion and antinociception) and thermoregulatory effects in mice and rats, consistent with the activity of nicotine itself (13, 16). Moreover, 6-MN was recently reported to be possibly more cytotoxic than nicotine in human bronchial epithelial cells (17). The receptor binding and functional potency of 6-MN at specified nAChR subtypes has not been determined. But this information is particularly important now because some manufacturers of vape solutions, e-cigarettes, and nicotine oral pouches reportedly add 6-MN (also known as metatine) to their products, including some that are advertised as “nicotine-free” (18, 19). Moreover, because 6-MN is not identical to nicotine and is found in only trace amounts in tobacco (20), it is not regulated by the U.S. FDA, although it is regulated in several other countries, including Australia, France, Germany, Belgium, and Sweden. Furthermore, because a significant fraction of vape solutions are thought to be purchased on-line through the gray market from manufacturers in countries where the regulation of 6-MN (and related compounds) is unknown, the possibility of high concentrations of 6-MN reaching users of vape solutions and oral pouches is further increased.

In this study, we quantitatively compared (-)6-MN and (-)nicotine with respect to receptor binding affinity and functional potency at α4β2 and α3β4 nAChRs, as well as potency to desensitize these receptors, which is one of the key effects of nicotine. In addition, because chronic administration of nicotine up-regulates the number of α4β2 nAChRs in rat (1, 21), mouse (2), and human brain (3, 4), we determined if 6-MN also does this in rat brain. The results across these two physiologically important nAChR subtypes reveal the potential scope and scale of 6-MN effects, which clearly indicate its high potency and the need for it to be regulated by the FDA or equivalent agency in the U.S.

## Results

### Binding of 6-MN and nicotine to α4β2 and α3β4 nAChRs

We compared the affinities of 6-MN and nicotine at human α4β2 and α3β4 nAChRs expressed in stably transfected HEK 293 cells in binding competition assays against 500 pM [^3^H]-EB. In parallel, we measured the affinities of these two drugs in rat cerebral cortex, where the predominant heteromeric nAChR is the α4β2 subtype (Flores et al., 1992) and in the rat pineal gland, where the predominant heteromeric nAChR is the α3β4 subtype (Hernandez et al, 2004). As shown in figures 1, both 6-MN and nicotine compete with high affinity for both nAChR subtypes expressed in HEK cells (**Fig. 1A and 1B**). Similarly, both drugs compete with high affinity for the two nAChR subtypes in native tissues (**Fig. 1C and 1D**). The results of these nAChR binding studies are summarized in Table 1. The binding affinities of 6-MN and nicotine are very similar at the α4β2 subtype heterologously expressed in our cells as well as in the rat cerebral cortex, which is not surprising given the structural similarity of the two compounds. In contrast, however, 6-MN binds with 5-times higher affinity than nicotine at α3β4 nAChRs expressed in our cells (p<0.01), and a similar 4-times higher affinity than nicotine at native α3β4 nAChRs in rat pineal gland (p<0.05).

**Table 1.** The binding affinities (K i values) were calculated from the IC 50 values for each tissue in Figure 1.

|  | K<sub>i</sub> (nM) |  |  |  |  |  |  |  |
| --- | --- | --- | --- | --- | --- | --- | --- | --- |
| | $\alpha 4\beta 2$ | | $\alpha 3\beta 4$ | | Rat Cortex | | Rat Pineal Gland | |
| 6-Methyl Nicotine | 4.0 | ± 0.4 | 49.2 | ± 7.0 | 9.2 | ± 0.02 | 78.8 | ± 21.1 |
| Nicotine | 4.5 | ± 0.6 | 262.6 | ± 28.2 | 7.3 | ± 1.0 | 307.0 | ± 71.9 |

**Figure 1.**
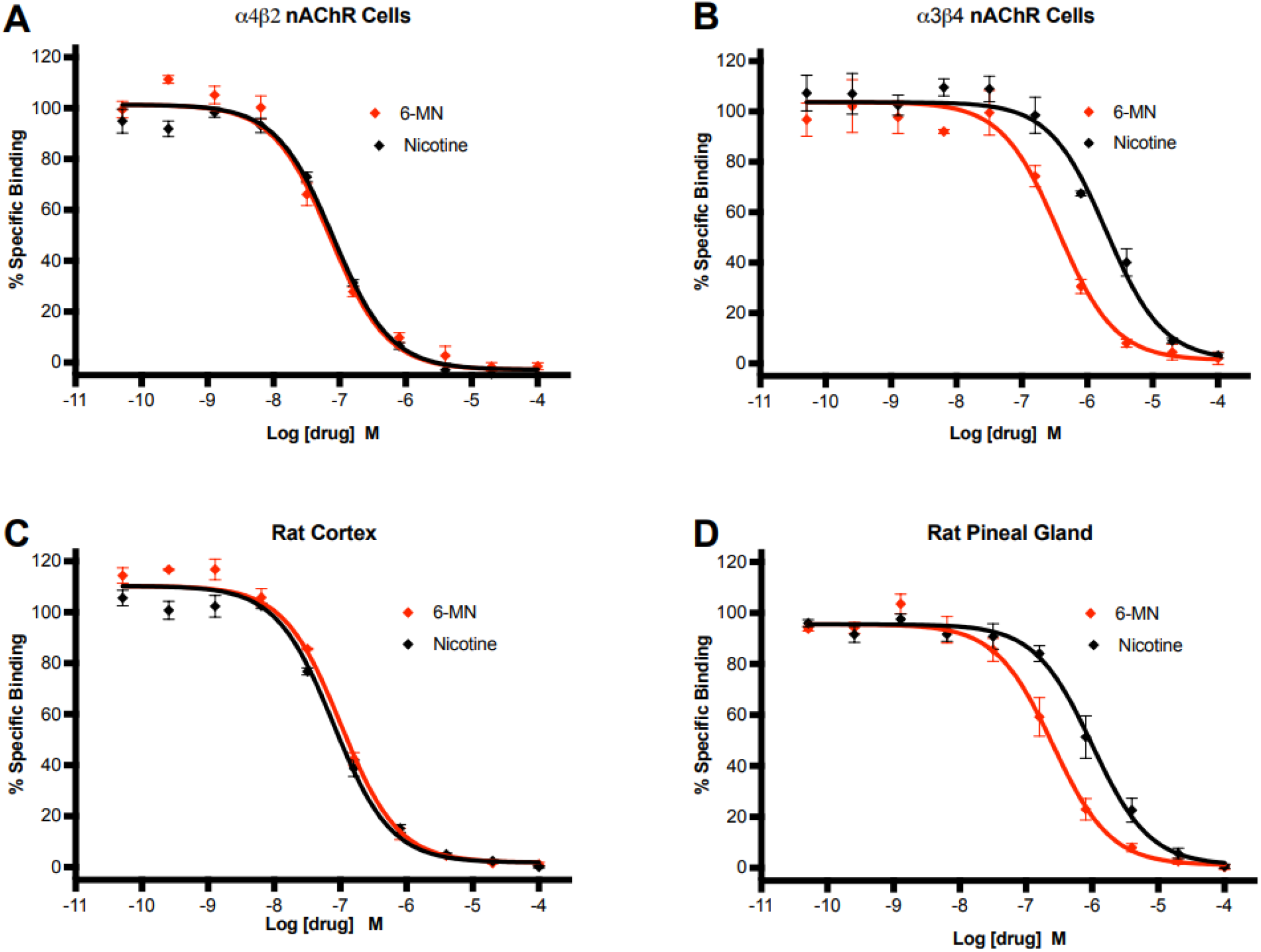
6-MN binds with high affinity to nAChRs. (A & B) 6-MN and nicotine compete effectively against 500 pM [^3^H]-epibatidine for binding to human α4β2 and α3β4 nAChRs stably expressed in HEK293 cells, and (C & D) in rat cerebral cortex and pineal gland, which express predominantly α4β2 and α3β4 nAChRs, respectively. See Table 1 for the binding affinities of 6-MN and nicotine at each of these nAChRs.

### Electrophysiology reveals activation and desensitization of α4β2 and α3β4 nAChRs by 6-MN

To compare the functional effects of 6-MN and nicotine on nAChRs, we performed whole-cell voltage-clamp recording in our nAChR-expressing HEK293 cells. Application of 30 µM 6-MN to α4β2 receptors evoked peak inward currents that were approximately 4-fold greater than the response to nicotine (**Fig. 2A & B**). Furthermore, 6-MN produced a much greater degree of acute desensitization of the current over the 10-s application period (**Fig. 2A & C**). The differences between 6-MN and nicotine were even more pronounced at α3β4 nAChRs; 6-MN stimulated currents approximately 10-fold greater than nicotine (**Fig. 2D & E;** note scale bars) and desensitized the response to a much greater degree (**Fig. 2D & F**). These results, in combination with our receptor binding data, indicate that 6-MN is more potent than nicotine at both of these nAChR subtypes, which is also borne out by the demonstrably slower deactivation rate of the current with 6-MN compared with nicotine.

**Figure 2.**
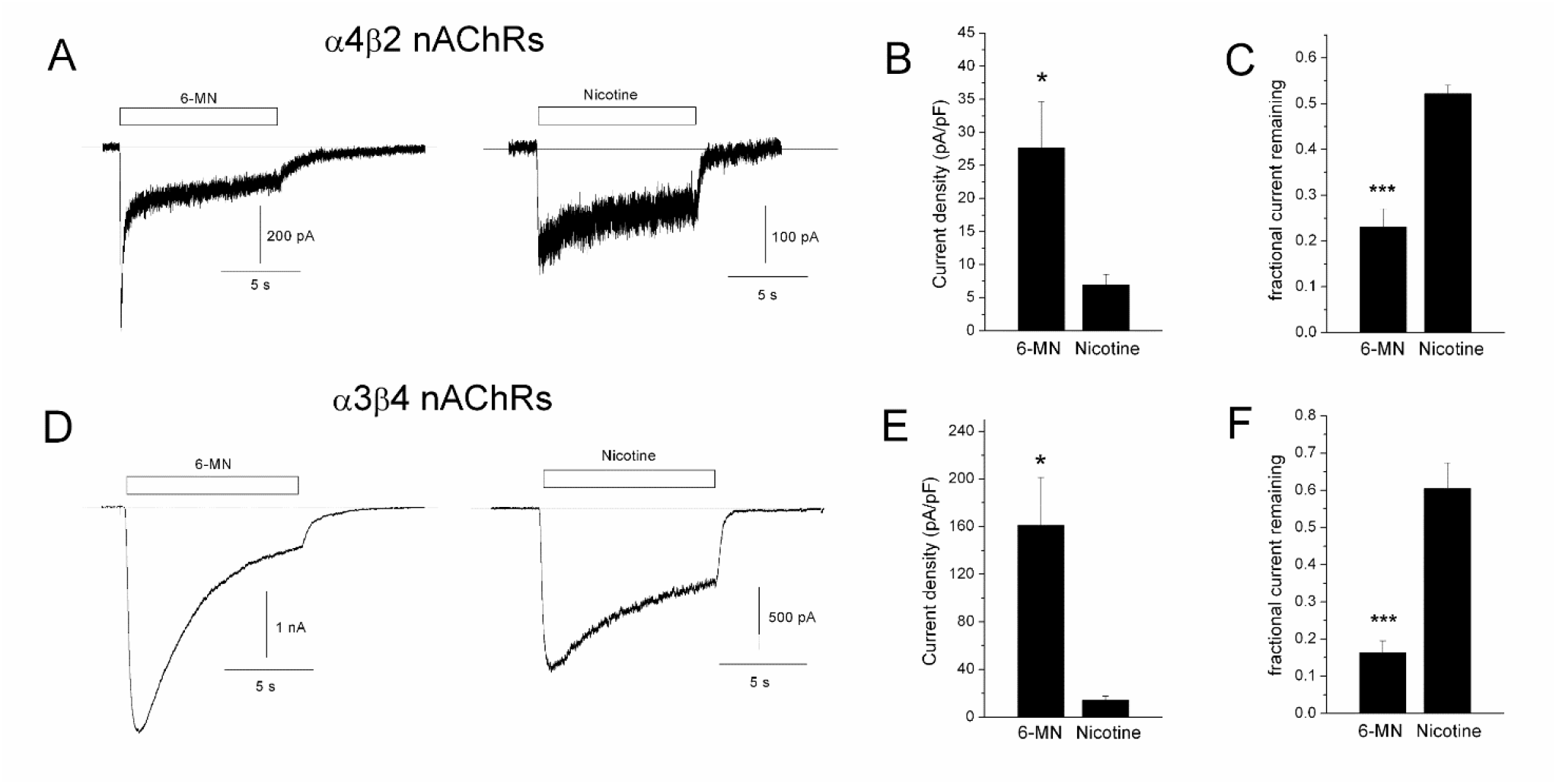
6-MN activates and desensitizes nAChRs measured in whole cell voltage-clamp recordings. (A & D) Representative inward current traces in HEK293 cells expressing α4β2 and α3β4 nAChRs in response to a 10 s application of 30 μM 6-MN or nicotine (note differences in scale bars). (B, C, E, & F) The mean current density and desensitization (final current divided by peak current) for 6-MN and nicotine evoked responses (α4β2 receptors, n=7 and 4; α3β4 receptors n=8 and 4; *p<0.05, ***p<0.001). The holding potential was -50 mV, and all data were obtained from the first drug application in separate cells.

### nAChR channel activation by 6-MN and nicotine measured in the aequorin bioluminescence assay

As a complementary approach and to quantitatively compare the agonist potency and efficacy of 6-MN and nicotine at nAChRs, we measured their activity in the aequorin bioluminescence assay. To do this, we first stably expressed the aequorin protein in our cells that also stably express the α4β2 and α3β4 nAChRs. When the cells are incubated with coelenterazine, it complexes with aequorin inside the cell; and when the receptors are subsequently activated, calcium enters the cell via the receptor channel and binds to the complex, which then emits light proportional to the calcium. The light signal is read and quantified immediately on a bioluminescence plate reader. Concentration-response curves for bioluminescence stimulated by 6-MN and nicotine at α4β2 and α3β4 nAChRs are shown in figure 3A & B. Analyses of these curves show that the two drugs stimulated a similar maximum response (E_max_) at both receptors, but the EC_50_ values indicate that at α4β2 nAChRs (**Fig. 3A**), 6-MN is 13-times more potent than nicotine (p<0.001), and at α3β4 receptors (**Fig. 3B**), it is 10-times more potent than nicotine (p<0.01). Moreover, the differences between 6-MN and nicotine are already evident at the lower concentrations that may be reached in brain and blood during vaping or use of a pouch, especially during initiation when adolescents may be first using these products.

**Figure 3.**
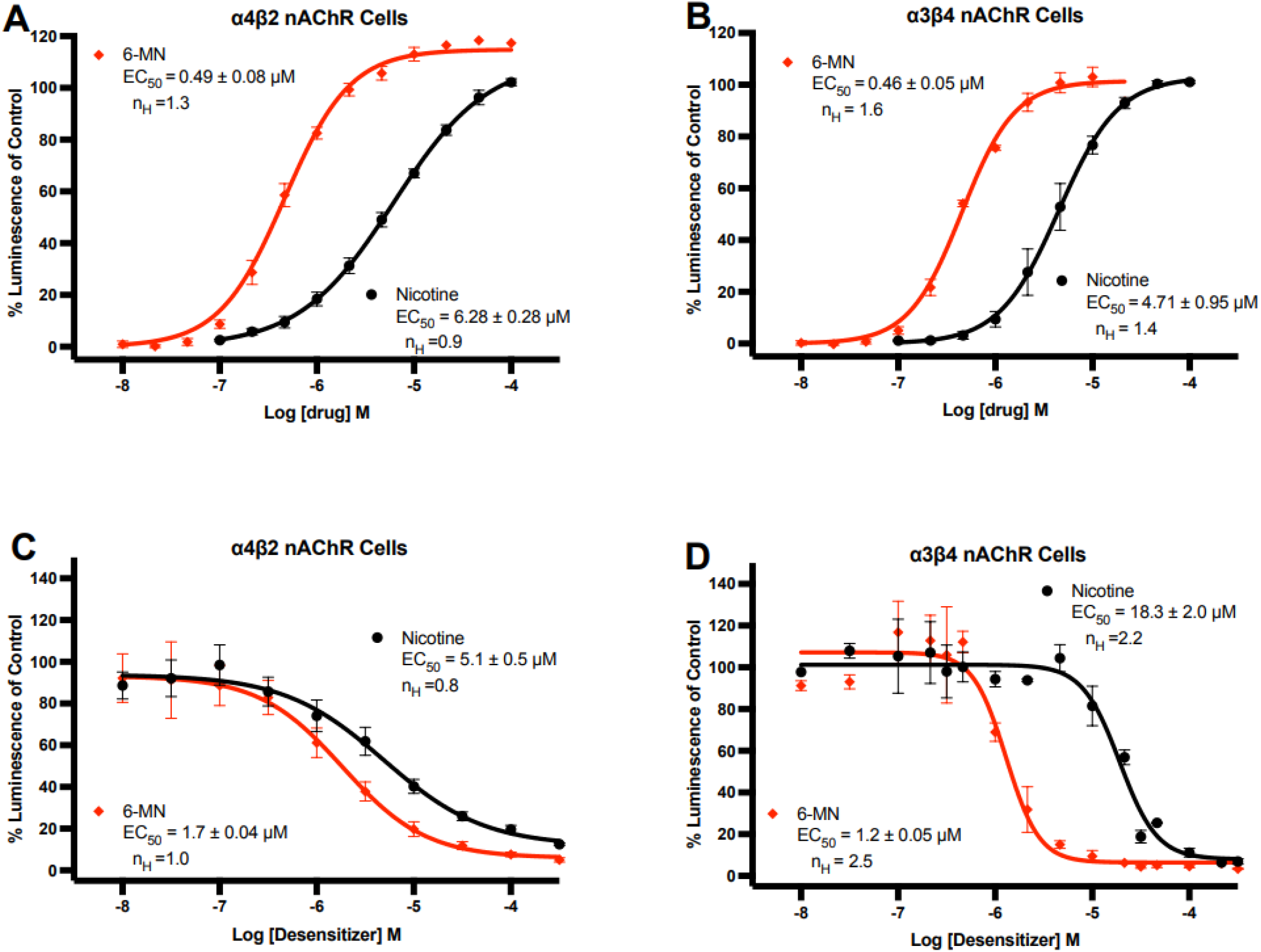
Both 6-MN and nicotine activate and desensitize nAChRs measured in aequorin bioluminescence assays. (A & B) 6-MN and nicotine activate α4β2 and α3β4 nAChRs expressed in HEK293 cells. Measured as the EC_50_ value for each drug, 6-MN is 13-fold more potent than nicotine at α4β2 receptors (n=3, p<0.001) and 10-fold more potent at α3β4 receptors (n=3, p<0.01). (C & D) 6-MN and nicotine desensitize α4β2 and α3β4 nAChRs. The response to 100 uM nicotine was measured after a two-minute exposure to each drug followed by a 10 min drug washout with fresh buffer.

### Desensitization of nAChRs by 6-MN and nicotine

To quantify and compare the desensitization of these nAChRs indicated in the electrophysiology studies, we measured the potencies of 6-MN and nicotine to desensitize α4β2 and α3β4 nAChRs in aequorin bioluminescence assays. For these studies, the cells were exposed for two minutes to 6-MN or nicotine at a concentration between 10 nM and 300 mM, washed for 10 min to remove the agonist, and then exposed to 100 mM nicotine, which in control cells stimulates a maximum bioluminescence response (see **Fig. 3A & B**). As shown in **figure 3C & D**, both drugs effectively desensitized the nAChR response, but as indicated by the EC_50_ values for desensitization after this 2 min exposure, 6-MN is 3-times more potent than nicotine at desensitizing the α4β2 receptor (p<0.01), and 15-times more potent than nicotine at desensitizing the α3β4 receptor (p<0.01). Moreover, while nicotine was ∼ 3.5-times more potent at desensitizing the α4β2 than the α3β4 receptor subtype (p<0.01), 6-MN was nearly equipotent at desensitizing the two receptor subtypes. Interestingly, the desensitization curves for both 6-MN and nicotine were much steeper at the α3β4 than at the α4β2 nAChRs, with Hill coefficients greater than 2 at the α3β4 receptors compared to 0.8 to 1.0 at the α4β2 receptors. This suggests that desensitization of the α3β4 nAChRs is complex and might involve positive cooperativity.

### Chronic administration of 6-MN increases α4β2 nAChRs in rat cerebral cortex

Chronic administration of nicotine increases (upregulates) nAChRs in rat, mouse, and human brain (1-4). The receptor increase appears to involve α4β2 nAChRs primarily or exclusively (21). This upregulation of nAChRs is thought to be a result of nicotine acting at a site different from the cell surface orthosteric binding site or the ion channel because it is not blocked by the competitive nAChR antagonist dihydro-β-erythroidine or the channel blocker mecamylamine (22). Instead, it is thought to involve an effect of nicotine in the endoplasmic reticulum within the neuron, probably by acting as a pharmacological chaperone to aid subunit assembly and maturation (23-25). To determine if chronic administration of 6-MN upregulates nAChRs in brain, we administered it to rats for 2 weeks at a nominal dose of 1 mg/kg/day via their drinking water, which also contained 2% saccharine. Control rats drank water containing 2% saccharine only. As shown in **Fig. 4**, this chronic treatment with 6-MN significantly increased nAChR binding in the cerebral cortex by about 25 percent (p<0.001), which is similar to the increases we’ve seen in rat cerebral cortex after chronic injections of nicotine (1, 22).

**Figure 4.**
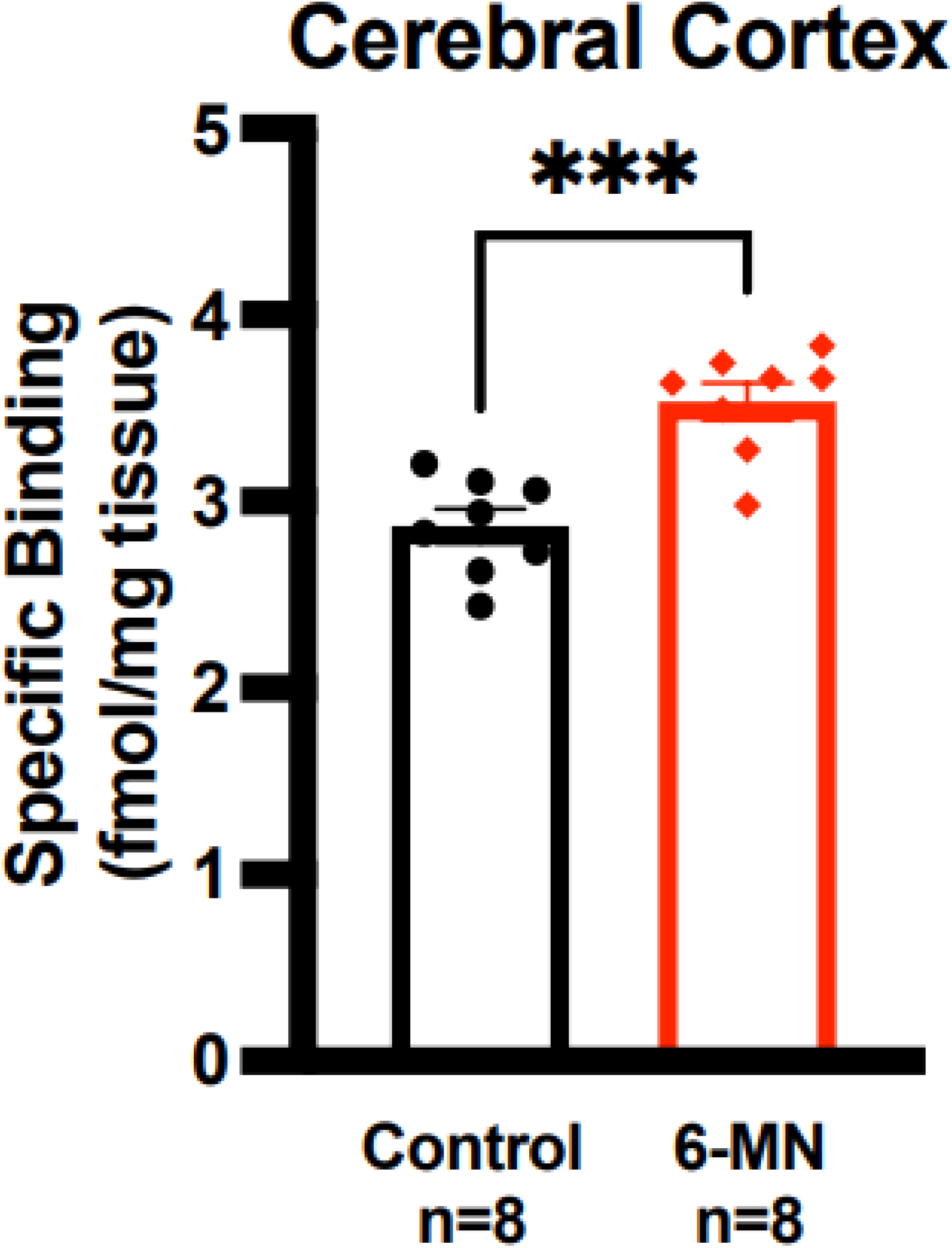
Chronic exposure to 6-MN increases (upregulates) nAChRs in rat brain. Male Sprague Dawley rats (∼300 g) were administered 6-MN at a nominal dose of 1mg/kg/day for two weeks in their drinking water, which also contained 2% saccharine to mask the bitter taste of the 6-MN. Control rats drank water with 2% saccharine only. Following sacrifice under isoflurane anesthesia, binding to nAChRs in cerebral cortex, which are predominantly α4β2 receptors, was measured with 500 pM [^3^H]-epibatidine. (n = 8 in each group; *** p<0.001).

## Discussion

The studies presented here demonstrate that 6-MN is a high affinity, high potency agonist at α4β2 and α3β4 nAChRs, two important nAChR subtypes in the nervous system. In fact, 6-MN is 13-fold more potent than nicotine in activating the α4β2 subtype of nAChR, which is the predominant nAChR in most areas of brain (21), and 10-fold more potent than nicotine in activating α3β4 nAChRs, which are found in certain areas of brain, and especially highly represented in the medial habenula and interpeduncular nucleus, which are implicated in the symptoms associated with withdrawal from nicotine (9, 11), as well as in brainstem nuclei associated with control of the autonomic nervous system (26). These receptors are also the major nAChR subtype in important autonomic nervous system ganglia (10), adrenal gland chromaffin cells (26, 27), and pineal gland (26, 28).

6-MN is 3-times more potent than nicotine in desensitizing α4β2 receptors, and 15-times more potent in desensitizing α3β4 receptors. The concentrations of 6-MN and nicotine to desensitize α4β2 and α3β4 nAChRs during their two-minute preincubation with the cells do not reveal an obvious relationship to either their affinities at the receptor binding sites or their potencies to activate the receptors. That may be due to the short preincubation period to induce desensitization. In previous studies, which used methods that allowed longer (60 min) pre-exposures, nicotinic agonists were 4-to10-times more potent to desensitize than to activate α3β4 receptors in rat PC12 cells (29) and in heterologous expression of α3β4 receptors (30).

Moreover, *in vivo* studies in rats indicate that after i.v. infusion, nicotine is 8-to 10-times more potent to desensitize than to activate the nAChR (likely the α4β2 subtype) that mediates prolactin release (31). Furthermore, after their brief activation by nicotine, the receptors remain desensitized for an hour or longer (32, 33) leading to the designation of nicotine as a time-averaged antagonist (31, 33). During vaping, of course, the exposure times to 6-MN and nicotine would be much longer, which would likely potentiate the degree and duration of desensitization of nAChRs at each concentration.

Based on its pharmacological characteristics described here, 6-MN administered via transdermal patch might be useful as a more potent nicotine replacement to help people trying to quit smoking cigarettes or vaping. Moreover, since the 6-postion of the pyridine ring of nicotine tolerates certain substitutions, and an analog compound with fluorine substituted at that position retains high affinity for the nAChRs in brain (Dukat et al., 1996), 6-[^18^F]nicotine could possibly be developed as an *in vivo* PET probe of these receptors.

At this time, however, the most important aspect of 6-MN is that it is added at unknown concentrations to some vape solutions, including some that are advertised as being nicotine-free, and because it is not structurally identical to nicotine and is not derived from tobacco, it is not regulated in the U.S., as it is in some countries. Unfortunately, this has opened the door to circumventing the main point of smoking and vaping regulations, which ostensibly is to prevent abuse of products that could maintain or even increase addiction to nicotine-like products. Furthermore, because 6-MN is more potent than nicotine at α4β2 and α3β4 nAChRs, it likely adds significantly to the addictive effects of nicotine in the brain in dual users of cigarettes and e-cigarettes. Moreover, when vaped, the highest 6-MN concentration would be in the lungs, where its high potency at α3β4 nAChRs would likely result in stimulation followed by desensitization of autonomic ganglia that influence the functions of the lung and heart, as well as more distal organ systems. The consequences of these actions are currently unknown.

The present findings contribute new and urgently-needed scientific data to inform the public and the FDA on the potential adverse health effects of current and future 6-MN products. If the FDA deems 6-MN-containing products as similar to tobacco products, like it did products containing synthetic nicotine, data from these studies would support adding 6-MN to the FDA’s list of harmful and potentially harmful constituents (HPHCs) in tobacco products, including vape solutions and pouches. This list is a critical regulatory mechanism, mandated by law, to identify tobacco constituents to surveil and, if feasible, regulate to protect public health (34). The present study lays the foundation for surveillance of other nicotine analogs as they emerge in new products.

Finally, the studies here demonstrate the concept that in addition to classifying a product with abuse potential or other health hazards based primarily on its chemical structure, an assessment of its pharmacological actions at the relevant receptors would be instructive.

## Methods

In all of the in vitro studies presented here (-)6-MN was compared with (-)nicotine.

### Cells

We used HEK293 cells stably expressing human α4β2 and α3β4 nAChRs. These cells with the rat and human versions of these nAChR subtypes have been described previously (Xiao and Kellar, 2004; Tuan et al., 2015).

### Rat tissues

Native α4β2 nAChRs were measured in cerebral cortex from male Sprague Dawley rats (∼250 g) purchased from Charles River Laboratories. Native α3β4 nAChRs were measured in rat pineal glands purchased in bulk from Zivik-Miller Laboratories.

### Binding affinities of 6-MN and nicotine at nAChRs

The affinities of 6-MN and nicotine for α4β2 and α3β4 nAChRs in our cell lines and in rat tissues were compared in receptor competition binding assays. Briefly, the cells are harvested in 50 mM Tris·HCl buffer (pH 7.4), washed in fresh buffer, homogenized with a Polytron homogenizer and centrifuged at 36,000 g. Aliquots of the resulting washed membranes in the pellet are resuspended in buffer and then incubated with 500 pM [^3^H]EB for 2 hr at room temperature in the presence of either 6-MN or nicotine at increasing concentrations (30 pM to 100 μM). Receptor-bound and free ligands are separated by vacuum filtration through Whatman GF/C filters treated with 0.5% polyethylenimine. The radioactivity retained on the filter is measured by liquid scintillation counting. Specific binding is defined as the difference between total binding and nonspecific binding assessed in the presence of 300 µM nicotine.

### Effects of 6-MN and nicotine on nAChR function. Electrophysiology

Whole-cell voltage-clamp recordings were performed by using an EPC8 patch-clamp amplifier (HEKA Electronik, Bellmore, NY) that was controlled by the program Pulse (version 8.65; HEKA Electronik). Data were collected at 5 KHz and low-pass filtered at 3 KHz. The bath solution contained 140 mM NaCl, 4 mM KCl, 1 mM MgCl2, 1.2 mM CaCl2, 10 mM Hepes, 5 mM glucose, pH 7.3. The pipette solution contained 140 mM CsCl, 10 mM HEPES, 5 mM EGTA, 1 mM MgCl2, pH 7.3. Solutions were applied via a valve-controlled gravity-fed perfusion system with an outlet (250-μm diameter) positioned ∼50 μm from the cell of interest. The solution exchange time was ∼200 ms.

### Aequorin bioluminescence assay

nAChR function was also evaluated with the aequorin bioluminescence assay, which was introduced to measure nAChRs by Stitzel and colleagues (Karadsheh et al., 2004). This assay is highly quantitative and provided assessments of the potency (EC_50_) and efficacy for activation, and EC_50_ for desensitization of 6-MN and nicotine at α4β2 and α3β4 nAChRs. For our assays, we first stably transfected the aequorin gene into our nAChR-expressing cells. When the cells are incubated with coelenterazine, it complexes with aequorin. When the receptor is subsequently activated, calcium enters the cell via the receptor channel and binds to the complex, which then emits light in proportion to the calcium. The light signal is read immediately and quantified on a bioluminescence plate reader (Promega). For assays to measure desensitization, cells were exposed to increasing concentrations of 6-MN or nicotine for two minutes and then washed for 10 minutes. The cells were then stimulated with 100 µM nicotine followed by buffer containing 100 mM calcium and 0.1% Triton X-100 (to permeabilize the cells), with immediate measurements of bioluminescence after each drug exposure. The bioluminescence evoked by 100 µM nicotine is defined as the 100% control value.

### In vivo studies of chronic 6-MN

To determine if chronic administration of 6-MN affects the density of nAChRs in rat brain, as nicotine does, male Sprague Dawley rats (∼300 g) were administered 6-MN for two weeks in their drinking water, which also contained 2% saccharine to mask the bitter taste of the 6-MN. Control rats drank water with 2% saccharine only. The concentration of 6-MN in the drinking water was based on rats drinking ∼10 ml/100 g body weight and intended to provide 6-MN at a nominal dose of 1mg/kg/day, which was chosen based on our experience using nicotine to upregulate nAChRs. At the end of the treatment period, the rats were sacrificed under light isoflurane anesthesia, the brain was removed and the cerebral cortex was dissected and kept frozen until used for receptor binding studies with [^3^H]-EB by procedures similar to those described above for the cells. All procedures involving animals were approved by the Georgetown University Animal Care and Use Committee and adhered to NIH ethical guidelines.

### Statistics

Differences between 6-MN and nicotine on effects of chronic administration of 6-MN on nAChR binding and function were compared by unpaired Student’s t-test. Data were analyzed and plotted using GraphPad Prism.11 software. An a level of p<0.05 was set as statistically significant.

## Data availability

The data that support the findings of this study are available on request from the corresponding author.

